# A representative Performance Assessment of Maximum Likelihood based Phylogenetic Inference Tools

**DOI:** 10.1101/2022.10.31.514545

**Authors:** Dimitri Höhler, Julia Haag, Alexey M. Kozlov, Alexandros Stamatakis

## Abstract

**Summary:** The evaluation of phylogenetic inference tools is commonly conducted on simulated and empirical sequence data alignments. An open question is how representative these alignments are with respect to those, commonly analyzed by users. Based upon the RAxMLGrove database, it is now possible to simulate DNA sequences based on more than 70, 000 representative RAxML and RAxML-NG tree inferences on empirical datasets conducted on the RAxML web servers. This allows to assess the phylogenetic tree inference accuracy of various inference tools based on realistic and representative simulated DNA alignments. We simulated 20, 000 MSAs based on representative datasets (in terms of signal strength) from RAxMLGrove, and used 5, 000 datasets from the TreeBASE database, to assess the inference accuracy of FastTree2, IQ-TREE2, and RAxML-NG. We find that on quantifiably difficult-to-analyze MSAs all of the analysed tools perform poorly, such that the quicker FastTree2, can constitute a viable alternative to infer trees. We also find, that there are substantial differences between accuracy results on simulated and empirical data, despite the fact that a substantial effort was undertaken to simulate sequences under as realistic as possible settings.

**Contact:** Dimitri Höhler

## Introduction

The field of phylogenetic inference deals with the problem of inferring hypotheses about the evolutionary relationships of species, mainly based on molecular sequence data. The inferred hypotheses are typically presented in the form of phylogenetic trees. There exist multiple approaches and optimality criteria to address this problem, such as distance based approaches (e.g., Neighbor Joining (NJ) (1)), Maximum Parsimony (MP) (2), Maximum Likelihood methods (ML) (3) or Bayesian inference methods (BI) (4–8). At present, the vast majority of phylogenetic analyses are conducted via ML and BI methods that both rely on the phylogenetic likelihood function that comprises an explicit stochastic model of sequence evolution. The inference of ML optimal trees is computationally hard (9) due to the super-exponential increase of possible unrooted binary tree topologies as a function of the number of taxa/sequences in the tree. Hence, there exists a plethora of inference tools which deploy different heuristic search strategies to find a tree with a *good* likelihood. Some of the most widely used tools are FastTree2 (10), IQ-TREE2 (11, 12), and RAxML-NG (13).

To assess the accuracy of the inferred trees and the performance of the respective heuristics, comparisons against alternative tools are typically based upon a combination of empirical and simulated datasets. The results of such performance assessments may deviate depending on the datasets used. To the best of our knowledge, there does not exist a standardized approach or set of benchmark data to assess and compare these tools. Instead, simulations and empirical datasets are conducted and selected predominantly in an ad hoc manner. Typical informal selection criteria for empirical datasets are the number of sites, the number of sequences (taxa), and the percentage of gaps or missing data in the datasets (see, for example, the respective papers describing FastTree2, IQ-TREE2, and RAxML-NG).

We examined the papers of the aforementioned analysis tools as well as a study comparing FastTree and RAxML (14) to determine the criteria deployed to select the respective benchmarking datasets. Then, we queried the RAxMLGrove database (15)(RG) for datasets satisfying these criteria. We found that at most 27% of datasets present in RG satisfy those criteria (see Table 1) and observe that the criteria do not appear to be representative if the data typically being analyzed.

**Table 1.** Criteria for selecting empirical (emp) and simulated (sim) DNA datasets analyzed in different publications and the respective proportion of dataset diversity in RAxMLGrove that they cover.

| Source | Selection criteria | Results | Representation |
| --- | --- | --- | --- |
| DNA RG | - | 73,786 | 100% |
| here: RG (sim) | < 470 taxa, < 18,764 patterns | 65,211 | 88% |
| IQ-TREE (11) | 200-800 taxa, #sites / #taxa $\leq 4$, #gaps $\leq 70\%$ | 8,339 | 11% |
| FastTree2 (10) (emp) | 500-239,882 taxa, 65-1,287 sites | 1,154 | 2% |
| FastTree2 (10) (sim) | 250-5,000 taxa | 8,091 | 11% |
| RAxML-NG (13) (emp) | 36-1,879 taxa, 18,328-37,350,521 sites | 4,391 | 6% |
| RAxML vs FastTree (14) (sim) | $\geq 1,000$ taxa | 1,471 | 2% |
| RAxML vs FastTree (14) (emp) | 117-27,643 taxa | 20,233 | 27% |

Related work assessing the inference accuracy of RAxML (16) and FastTree (17) has been conducted by Liu *et al*. (14). The authors used 1, 800 simulated and 10 empirical datasets and concluded, that RAxML (the predecessor of the re-designed code RAxML-NG that implements essentially the same algorithm) generally outperforms FastTree in terms of topological accuracy on smaller datasets, but that the differences diminish on large datasets with respect to the number of taxa. We make a similar observation for the updated versions of the two ML tools based upon around 5, 000 empirical datasets from TreeBase. Further, we can characterize and classify these datasets in a systematic manner by predicting their difficulty scores via the Pythia (18) tool. Pythia assigns values between 0 (easy/strong signal) and 1 (difficult/weak signal). Here, difficulty essentially quantifies the signal strength in the data. Zhou *et al*. (19) perform a large-scale analysis of ML-based methods on 19 empirical datasets with up to 200 taxa and partially thousands of genes. They find, that RAxML and IQ-TREE perform similarly in terms of topological accuracy on single-gene datasets, while FastTree performs substantially worse. In our study, we confirm this result for empirical datasets (composed of single- and ca. 10% multi-gene data) with a difficulty score below 0.7.

The key contribution of our work is that we deploy empirical and simulated DNA data that representatively covers at least 75% of the data in the two databases (TreeBase and RAxMLGrove). We assess accuracy by means of the inferred tree topologies, the ML scores, and their statistical plausibility (i.e., whether the trees differ significantly from the best-known tree).

We did not include amino-acid (AA) datasets here to reduce the computational burden and CO_2_ footprint of our analyses. We conducted our experiments using a Snakemake (20) pipeline, which is available at https://github.com/angtft/PhyloSmew. This pipeline can be easily extended by additional phylogenetic inference tools and will thus contribute to conducting accuracy inference studies in a more standardized way.

## Materials and Methods

In this section, we first briefly describe the RAxMLGrove database (15) that we use for simulating Multiple Sequence Alignments (MSAs). Then, we describe the methods for selecting and simulating datasets. We undertake an effort to create realistic simulated MSAs, especially in terms of realistic gap patterns, and we evaluate this attempt in the respective Section on simulation quality. Then, we briefly describe our selection strategy of empirical datasets from the TreeBASE database (21).

### RAxMLGrove Database

Throughout our experiments, we extensively use the RAxMLGrove (RG) database. RG contains anonymized phylogenetic trees inferred by users of RAxML/RAxML-NG web servers at the San Diego Super-computer Center (22) and the SIB Swiss Institute of Bioinformatics (https://raxml-ng.vital-it.ch) with their respective inferred model parameters and program execution logs. At present, RG comprises more than 70, 000 datasets and is continously growing. The RAxMLGroveScript repository (https://github.com/angtft/RAxMLGroveScripts) provides functionality for simple data access using an SQLite-database with a corresponding Python script (henceforth called RGS). In addition to dataset queries based on a set of dataset characteristics (e.g., the number of taxa, or branch length variance of the tree), RGS provides the `generate` option to download datasets and generate simulated MSAs based on the estimated model parameters of that dataset. The simulations are conducted by executing Dawg (23) or AliSim (24). Since simulation is the main RGS functionality of interest in this work, we briefly list and describe the contents of one such downloaded simulated dataset, after using the `generate` command, in Table 2.

**Table 2.**
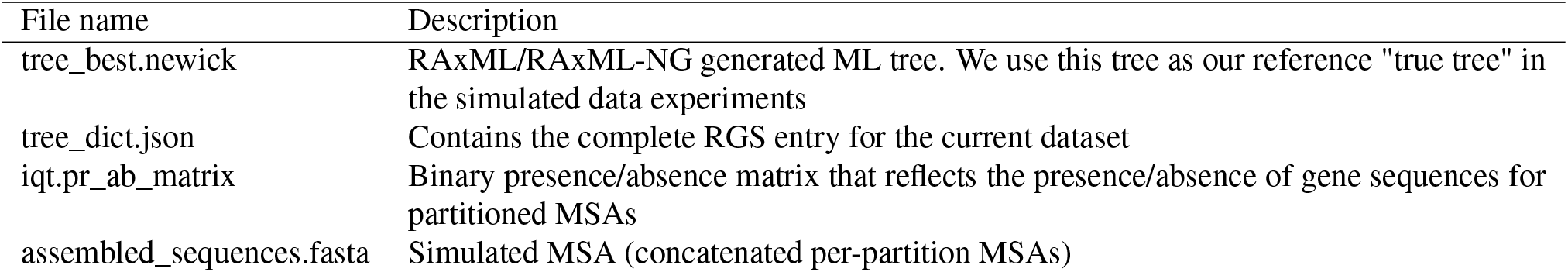
Files created by the RGS command *generate*

### Simulated Data

We generate simulated data based on the RG database, version 0.7. As our goal is to assess ML tool inference behavior for *representative* datasets, we proceeded as follows to select the datasets for our analysis:

1. For the sake of simplicity, we only selected trees inferred on DNA alignments under the General Time Reversible (GTR) (25) model.
2. We selected datasets from the 95-percentiles of the number of taxa and number of site patterns (i.e., the numbers of unique sites in an MSA) distributions respectively, that is, datasets with a number of taxa *<* 470 and a number of patterns *<* 18, 764. Thereby, we omitted computations on excessively large datasets, which can be considered as outliers.
3. We initially sorted the datasets by their site patterns per taxa ratio and subsequently divided them into 20, 000 buckets. From every bucket, we randomly selected one dataset as a *representative* of that bucket.
4. For every representative, we simulated one MSAs using the corresponding RGS functionality, which we describe in more detail below.

Under these criteria, we selected 20, 000 DNA datasets representing more than 80% of overall RG data in terms of signal strength (approximated by patterns/taxa ratio) and simulated MSAs based on the inferred ML trees and their respective inferred model parameters.

During the simulation process, we encountered the problem of simulating MSAs with gaps. To simulate MSA gaps in a realistic way, the RG entry ‘OVERALL_GAPS’ that reflects the number of gaps in the original empirical MSA is not sufficient. According to Haag *et al*. (18), the fraction of gaps in an MSA, does not have a substantial impact on dataset difficulty. Thus, one possibility is to avoid gaps in simulations and to solely execute the tests on MSAs without gaps. However, this induces another problem. While the mere gap fraction does not constitute a reliable difficulty predictor, the specific occurrence pattern of simulated gaps via an appropriate simulated insertion/deletion process affects the number of distinct site patterns in the MSA. In turn, this affects the resulting difficulty, as the difficulty *is* correlated with the patterns/taxa ratio (see Figure 1). In our initial experiments, we observed differences exceeding 20% between the number of site patterns in the simulated MSAs *without* an insertion/deletion process and the number of patterns in the respective RG entries.

**Fig. 1.**
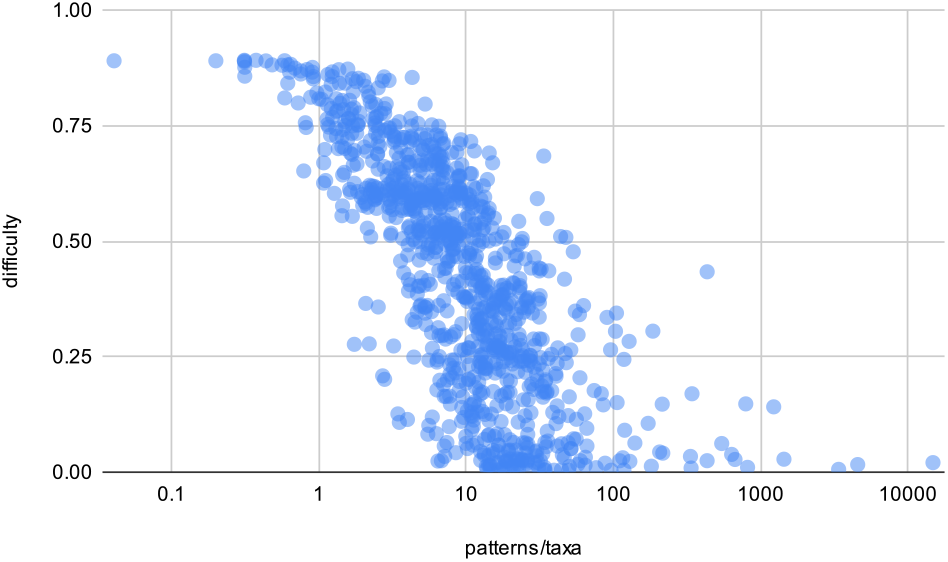
Predicted TreeBASE MSA difficulty with Pythia of 1,000 randomly selected datasets in correlation to the patterns/taxa rate (logarithmic x-axis).

We used AliSim (24) to simulate MSAs for our experiments. For simulating gaps, AliSim offers the `--indel` and `--indel-size` parameters. The `--indel` parameter is a user-specified rate for generating insertions and deletions (so-called *indels*) at sites during the sequence simulation process. The indel lengths are drawn from distributions, which can be specified using the `--indel-size` parameter. Currently, AliSim supports the Geometric, Negative Binomial (NB), Zipfian, and Lavalette distributions originally proposed in (26), which can be separately set and selected for the insertion and deletion process. Different distributions also need to be parameterized differently (e.g., mean and variance for indel sizes for the Negative Binomial distribution). Thus, even if we use the same distribution for insertions and deletions, we need to set a total of 6 parameters, that is, the insertion and deletion rates as well as the respective means and variances of the insertion and deletion sizes to parameterize the NB distribution. An additional complication is that the sequence length parameter used by some simulators (e.g., AliSim) only defines the sequence length of the single starting sequence on which the simulation is then carried out along the tree. Thus, when gaps are introduced during the simulation process, the overall MSA length will typically exceed the seed/root sequence length.

Under the assumption, that a simulation under the ‘correct’ settings for these 6 parameters will result in a simulated MSA where the differences between the number of sites, number of patterns, and gap percentage of RG entries and simulated MSAs are minimal, we can define an evaluation function *d* to quantify how well the simulated MSA matches the original MSA. Let *s*_*orig*_, *p*_*orig*_, *g*_*orig*_ and *s*_*sim*_, *p*_*sim*_, *g*_*sim*_ be the number of sites, number of patterns, and fraction of gaps in the RG entries and the simulated MSAs, respectively. Further, let *w*_1_, *w*_2_, *w*_3_ be arbitrary, yet constant weights. We can then define *d* as follows:

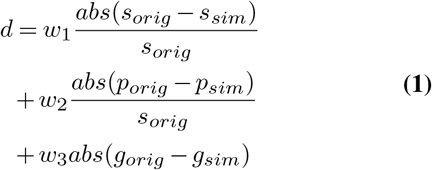

Since it is unclear, how to set these parameters such as to minimize *d* for a given simulated dataset, we resorted to Bayesian optimization (27–29), using the Python library skopt (30), and implemented the Bayesian Optimized iNdel seeKer (BONK) in RGS. Given *w*_1_, *w*_2_, *w*_3_, a scaling factor *c*_*seq*_, insertion and deletion size distributions, and a maximum number of optimization rounds *n*_*opt*_, BONK uses the optimizer to iteratively explore the indel rates and size distributions to minimize *d* for the simulated MSA. For the sake of simplicity, we arbitrarily chose (preliminary tests indicated no apparent reason to choose one distribution over another) the NB distribution for both, insertion, and deletion sizes. We use *c*_*seq*_ to define the sequence length range for the root sequence. As stated above, the length of the root sequence has to be chosen as a function of the gaps being inserted to minimize the difference between *s*_*orig*_ and *s*_*sim*_. The length interval is defined as [*s*_*orig*_ × (1 − *c*_*sim*_), *s*_*orig*_ × (1 + *c*_*sim*_)]. Preliminary experiments have also shown that, apart from *n*_*opt*_, the minimization of *d* is strongly affected by the values of *w*_1_, *w*_2_, *w*_3_. These weights penalize some property differences more than others, and since the number of sites, number of patterns, and gaps are correlated, it appears that choosing these weights is not trivial.

Thus, we deployed a second optimizer (results not shown) to optimize the weights with respect to the average *d* of simulations based on random subsets of datasets from RG. We found that *w*_1_ = 9, *w*_2_ = 10, *w*_3_ = 1, *c*_*seq*_ = 0.7 worked sufficiently well, and used these weights for all our simulations with BONK.

After implementing BONK, we discovered SpartaABC (31, 32), a tool which uses the Approximate Bayesian Computation method (33, 34) to estimate the indel rates and sizes of a given MSA. The authors use a far more sophisticated distance function than *d*, overall comprising 27 features. Since the original MSAs are not available in RG, we cannot compute their proposed distance function and include SpartaABC into RGS. Thus, our simulations were conducted with BONK. However, we followed the suggestions regarding SpartaABC (and references to empirical studies) and made adjustments to BONK: We set the explored indel intervals to [0.0, 0.05], switched to the Zipfian distribution for indel sizes, and set the explored interval of the corresponding *a* shape parameter to [1.001, 2].

In addition to the aforementioned procedures, we also use the presence/absence matrices for partitioned datasets, when available in RG. A presence/absence matrix is a 2-dimensional binary matrix denoting the presence of information (sequence data) for taxon *k* at partition *l*. If there is no sequence data for taxon *k* at partition *l* we set *M* [*k, l*] := 0, and *M* [*k, l*] := 1 otherwise. After the simulation process, we therefore remove per-partition sequences from the MSA according to *M*. This is conducted before calculating *d*, such that the optimizer is aware of the gaps introduced by *M*.

### Simulation Quality

We evaluated our BONK method for gap-aware simulations by comparing the absolute differences between the properties (i.e., number of sites, number of site patterns, and gap proportion) of the simulated MSA and the respective RG database entry. We show the normalized differences in Table 3. In the worst case, that is, when the optimizer is not able to find parameters improving upon trivial simulations, we obtain an MSA without gaps. On average, we are able to simulate MSAs with properties which are closer to those of the RG entries than under trivial simulations without an indel model in terms of pattern numbers. However, due to the weights in our distance function, the difference in the gap proportions remained roughly the same - with the difference, that we now often insert more gaps than necessary.

**Table 3.**
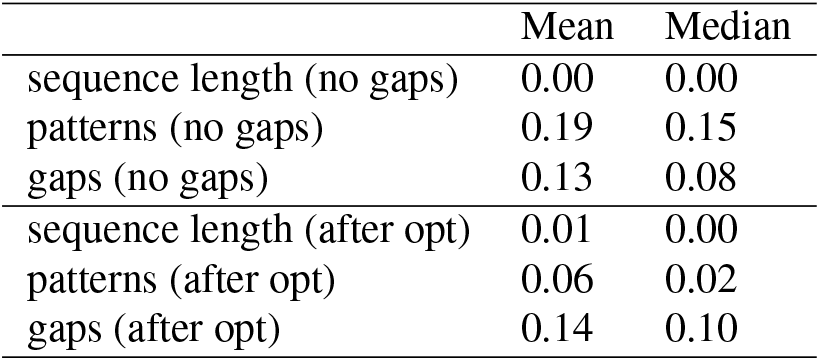
Normalized (absolute) MSA property differences between RG entries and simulated MSAs. We compare the trivial (no gap insertion) with the optimized simulation using BONK.

|  | Mean | Median |
| --- | --- | --- |
| sequence length (no gaps) | 0.00 | 0.00 |
| patterns (no gaps) | 0.19 | 0.15 |
| gaps (no gaps) | 0.13 | 0.08 |
| sequence length (after opt) | 0.01 | 0.00 |
| patterns (after opt) | 0.06 | 0.02 |
| gaps (after opt) | 0.14 | 0.10 |

We further compared the difficulty of the simulated and original MSAs using Pythia (18). For this, we randomly selected a subset of 989 TreeBASE DNA datasets (note that the original empirical MSAs are required for difficulty prediction, which are available in TreeBASE but not in RG), using the same size-based dataset selection criteria as in the preceding experiments, inferred a RAxML-NG tree using 50 parsimony and 50 random starting trees under the GTR+Γ model and used the inferred best-found tree, and model, to conduct the following distinct MSA simulations: (1) AliSim simulation without any indels; (2) AliSim simulation with the so-called mimicking function, which uses a specifiable (original) MSA to first infer a tree, then simulate an MSA, and subsequently superimpose the gap pattern from the original MSA to the simulated one; (3) BONK; (4) the SpartaABC tool, albeit with modifications to its internal pipeline. In SpartaABC, we removed the realignment step and reduced numbers of default burn-in and optimization rounds (1, 000/10, 000 instead of 10, 000/100, 000 respectively) to reduce the overall computational time, as some of the MSAs required more than 24 hours for a single indel rate inference. After the analysis, we used the suggested indel parameters by SpartaABC to simulate 10 MSAs with AliSim and selected the MSA with the lowest distance (as defined in SpartaABC) to the original MSA. We are fully aware that we do not fully utilize the capabilities of the SpartaABC algorithm, but we include it here to provide an intuition of how this method could perform in terms of difficulty matching, since it might be a valuable addition to the RGS simulation procedure.

In Table 4, we observe the following: Overall, there are small absolute differences between (1), (3), and (4). One might argue that the differences of these three approaches fall within the margin of error of the difficulty prediction. Surprisingly, inserting no gaps at all yields the difficulty prediction that is closest to the original difficulty, albeit the number of patterns more closely matches the original when using (3) instead of (1), and the gap proportion errors of (1) and (3) with respect to the original are similar. (1) and (3) do not require additional information about the underlying MSA, other than the properties already available in RG. Method (3) requires *n*_*opt*_ = 100 steps, which means that it is at least 100 times slower than the single MSA simulation conducted by method (1). Method (4) requires an a priori computation of the gap features for the SpartaABC distance function. This could potentially be included into a future RG release, as it requires little additional memory to store these data and does not expose the original MSA to the public (i.e., it can directly be computed on the respective web-servers before being stored in RG). Method (2) is probably the most intuitive approach. Here, the distance *d* is negligible on average (data not shown). This method performs worst in terms of difficulty differences. We believe that different sites can have different importance for the inference (this seems at least to be true for empirical data (35)), which a gap insertion by superimposing gaps does not take into account. Method (2) also requires the complete original MSA (if using the default AliSim mimick function) or at least the gap matrix of the original MSA to be available (if using the already inferred best tree and model present in RG).

**Table 4.**
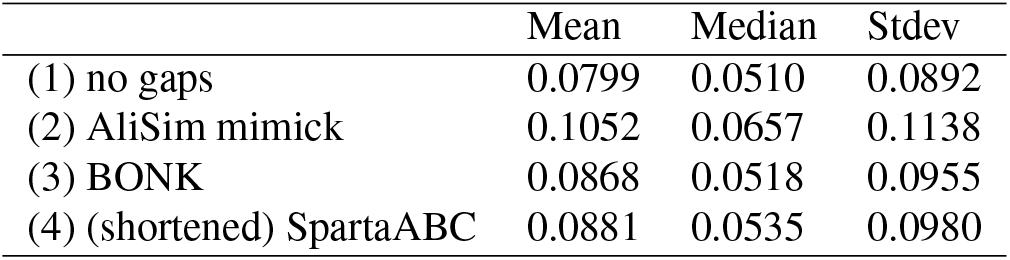
Absolute differences in difficulty between the original (empirical) MSAs and the simulated MSAs (based on 1000 datasets from TreeBASE, selected from 95-percentiles of taxa and site pattern numbers).

|  | Mean | Median | Stdev |
| --- | --- | --- | --- |
| (1) no gaps | 0.0799 | 0.0510 | 0.0892 |
| (2) AliSim mimick | 0.1052 | 0.0657 | 0.1138 |
| (3) BONK | 0.0868 | 0.0518 | 0.0955 |
| (4) (shortened) SpartaABC | 0.0881 | 0.0535 | 0.0980 |

### Empirical Data

We conducted experiments on 5, 000 MSAs selected from TreeBASE (21). The dataset filtering criteria and the subsequent selection of representative datasets are analogous to the criteria for selecting simulated data from RG: We selected datasets from the 95-percentiles of number of taxa and site patterns, that is, datasets with a number of taxa *<* 214 and a number of site patterns *<* 3, 475. Multiple datasets contained sequences which were completely empty/undetermined. As these sequences cannot be placed onto a tree in any meaningful way, and the ML inference tools handle completely undetermined sequences differently (e.g., IQ-TREE2 terminates instantly, RAxML-NG does not, but inserts such sequences at random), we excluded datasets containing completely undetermined sequences from the analyses. During the experiments, 4 datasets containing ambiguous character encodings were excluded, since IQ-TREE2 incorrectly attempts to analyze them as AA datasets (if our interpretation of the log file is correct) and terminates with an error (this issue has already been reported by a user in the IQ-TREE Google Group). Furthermore, 2 datasets were excluded as they triggered assertions in RAxML-NG to fail during the tree search process. Overall, more than 75% of the datasets diversity (in terms of pattern/taxa ratio) in Tree-BASE were covered.

### Evaluation Pipeline

We implemented our experimental workflow via a Snakemake (20) pipeline. The datasets were selected, prepared, and generated as described above. The main steps of the pipeline are the following (also, see Figure 2):

1. Query the database for datasets, sort the results based on the number of taxa by the number of site patterns ratio, and split the results into *n* buckets
2. Randomly pick a dataset from each bucket
3. Run the FastTree2, IQ-TREE2, and RAxML-NG inferences under the General Time Reversible (GTR) substitution model in combination with the Γ model of rate heterogeneity. To obtain parsimony trees, select the parsimony trees with the highest log likelihood (LnL) score from the set of inferred RAxML-NG parsimony starting trees. Then, use the tree evaluation function of RAxML-NG (which optimizes branch lengths and model parameters while keeping the tree fixed) on all inferred trees to obtain a consistent and comparable LnL score
4. Compute pairwise Robinson-Foulds distances (36) and LnL-differences between the true trees or best-known ML trees (on empirical data) and the inferred ML trees and run the topological significance tests of IQ-TREE2 on the complete set of trees (including the true tree)
5. Plot accuracy statistics

**Fig. 2.**
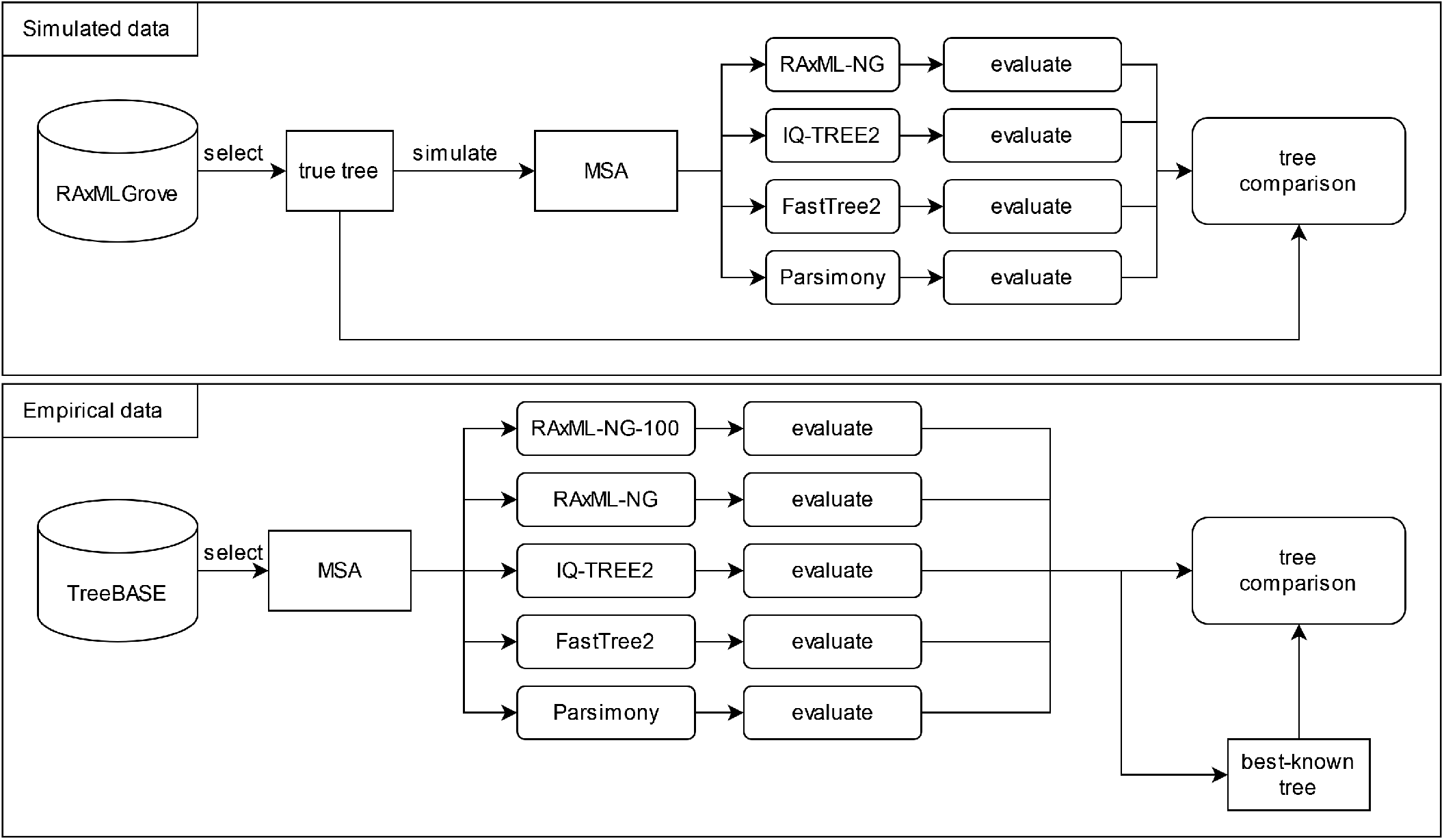
Flowchart of the conducted experiments on simulated and empirical data.

For the tree inferences, we used the respective default parameters to reduce the complexity of the experimental setup:

#### RAxML-NG

~~~
raxml-ng --msa assembled_sequences.fasta
 --model GTR+G --prefix [prefix]
 --seed [seed] --threads 4
 --force perf_threads
~~~

For partitioned datasets we substituted “–model GTR+G” with “–model [partition_file]” as all original partition files were available for empirical datasets. For simulated datasets, RG contains per-partition substitution model and MSA length parameters, based on which RGS simulates the MSAs and generates the partition files.

#### IQ-TREE2

~~~
iqtree2 -s assembled_sequences.fasta
 -m GTR+G -nt 4
 --prefix [prefix]
~~~

For partitioned datasets, we appended “-p [partition_file]”.

#### FastTree2

~~~
FastTreeDbl -gtr -gamma
 -nt assembled_sequences.fasta
~~~

To the best of our knowledge, FastTree2 does not support partitioned analyses and therefore all analyses were conducted on unpartitioned MSAs. We could not evaluate some of the FastTree2 trees via RAxML-NG as they contained multifurcations. In that case, we resolved the multifurcations at random prior to LnL evaluation.

## Results and Discussion

We evaluated the tree inference accuracy via the Robinson-Foulds (RF) distance (36) (as implemented in RAxML-NG) as well as Quartet-distances (37) (implemented in tqDist (38)), and log-likelihood (LnL) differences. We further applied statistical significance tests (as implemented in IQ-TREE2) using the RELL method (39), including Bootstrap Proportion, Expected Likelihood Weights (40), and the Approximately Unbiased Test (41). We used the topological distances and LnL score comparisons to compare the true (simulated data) or best-known (empirical data) ML trees with the inferred trees. In absence of a true tree for the empirical datasets, we executed a more thorough RAxML-NG search (100 independent tree searches, using 50 parsimony and 50 random starting trees) to find the tree with the best-known LnL score. If one of the other inference tools inferred a tree with a higher LnL, we re-defined that tree as the “true” tree with the best-known LnL score. In the statistical tests, we compared the number of times the inferred trees passed the tests with 95% confidence (compared to the best-known tree). We will refer to trees passing at least one test as *plausible* trees. If a tree *t*_1_ passes more tests than tree *t*_2_, we refer to *t*_1_ as being *more plausible* than *t*_2_.

After the analysis, we divided the results into 5 buckets based on the estimated MSA tree inference difficulty computed by Pythia (18).

Figure 4 shows the results of these experiments on empirical data. We excluded Quartet distances from the plots for the sake of readability, as they do not provide any additional insights (see Supplement for plots with quartet distances). Generally, there is a trend for inference accuracy to decrease and become more variable (spread-wise) with increasing difficulty. This also shows that the difficulty measure is meaningful in practice. In terms of LnL scores, RAxML-NG finds the trees with best LnL scores on average at all difficulty levels, IQ-TREE finds the second-highest scores, and parsimony and FastTree2 yield trees with substantially lower scores. The ML scores, as evaluated by RAxML-NG, of parsimony- and FastTree2-based inferences tend to be relatively close to each other. For MSAs exhibiting low (*<* 0.2) and high (*>* 0.8) difficulty levels, the absolute LnL differences to the best-known tree in the majority of our experiments are in the single digit (log likelihood units) range (see Figure 3).

**Fig. 3.**
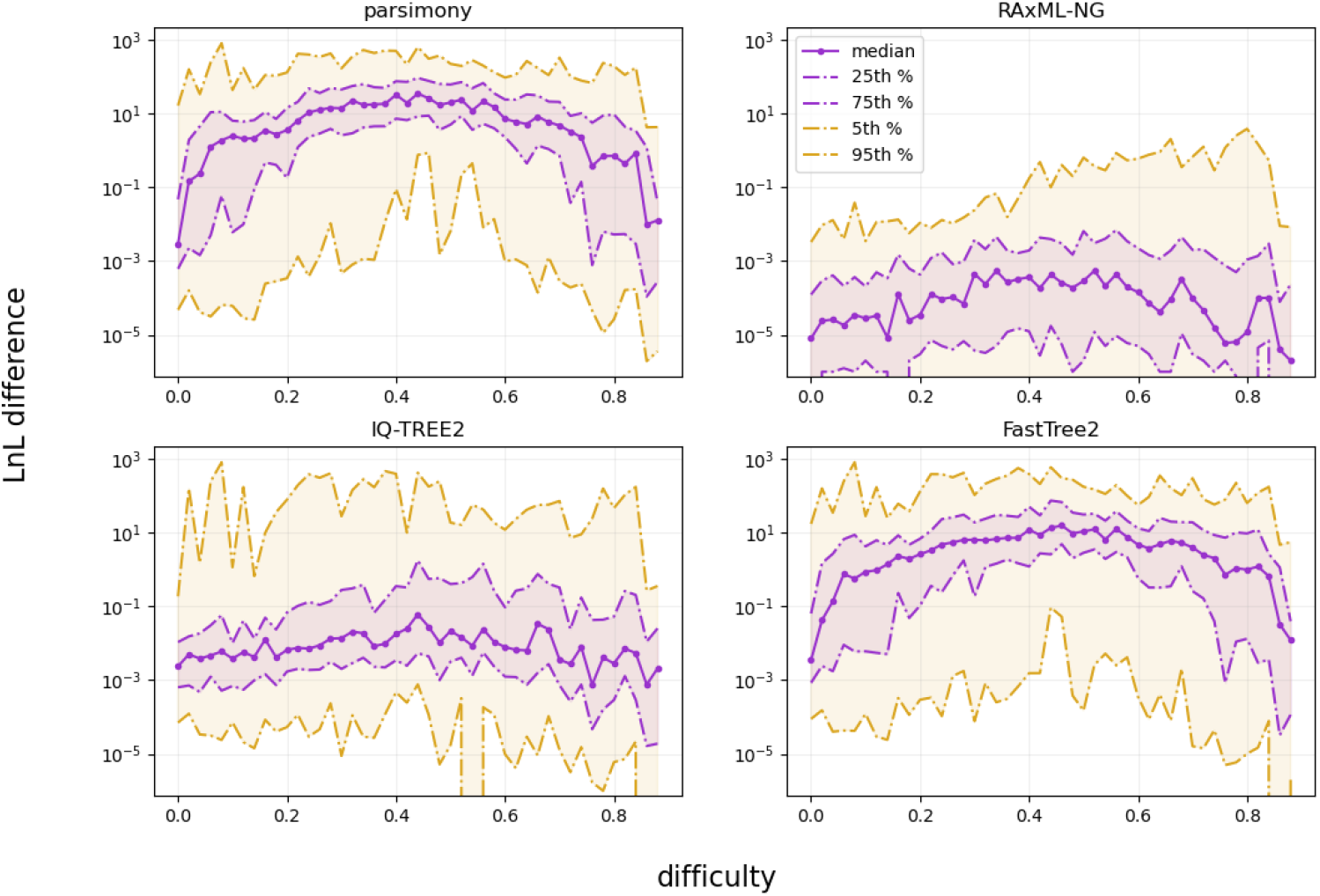
Absolute log-likelihood (LnL) score differences (log scale) from the best-known ML tree on TreeBASE data.

**Fig. 4.**
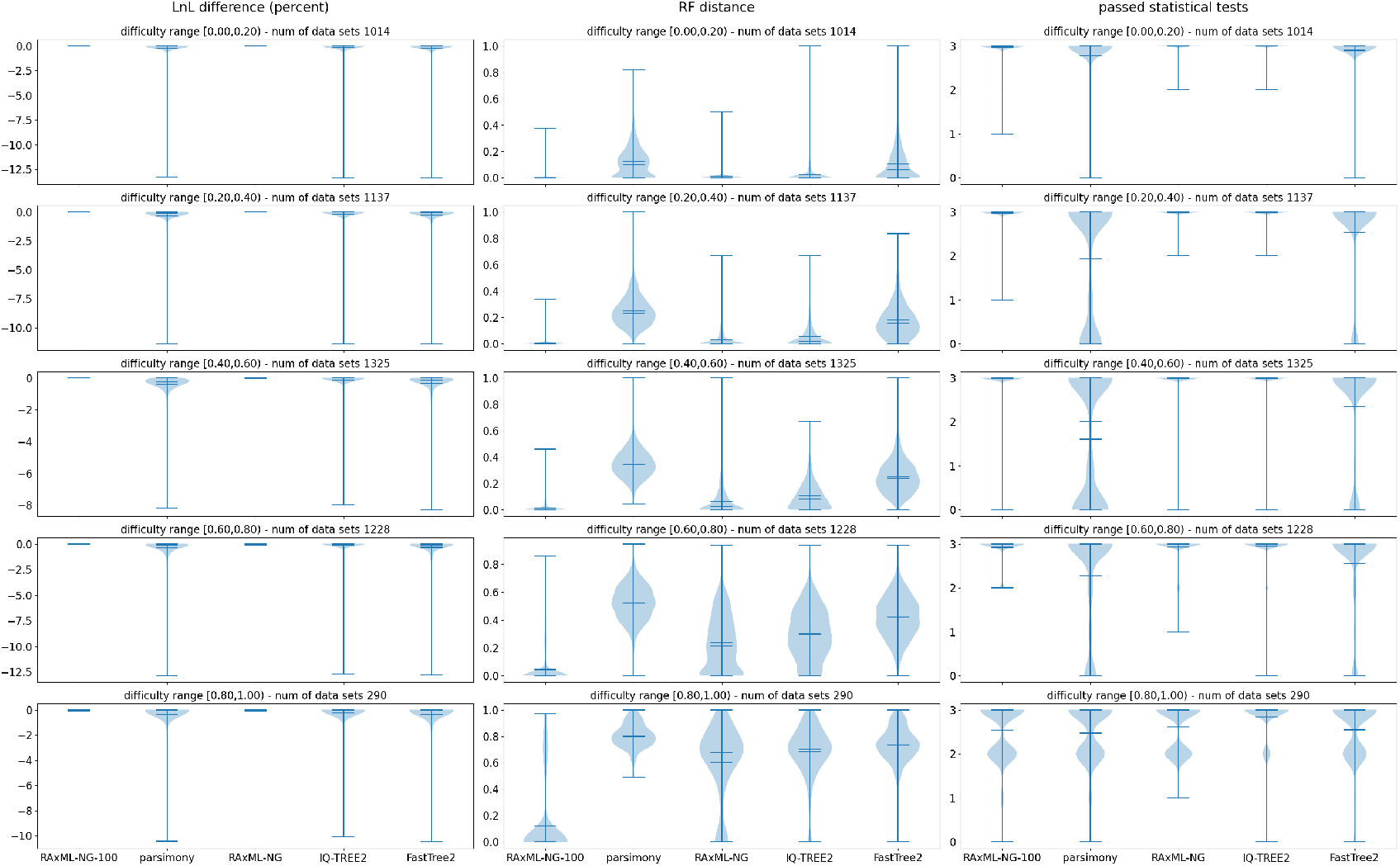
Relative log-likelihood (LnL) score differences, and RF-distances to the best-known ML tree, and numbers of passed statistical tests of all inferred trees on empirical TreeBASE datasets. “RAxML-NG-100” denotes the tree inferred by conducting 100 tree searches using RAxML-NG.

We observe a trend of increasing RF-distances for every tool with increasing difficulty. On average, RAxML-NG finds the tree that is closest to the true or best-known tree on datasets with a difficulty below 0.8. IQ-TREE2 is a close second. On MSAs exhibiting a high degree of difficulty, the differences between all analyzed tools appear to be negligible.

With respect to the statistical tests, IQ-TREE2 consistently finds the largest number of plausible trees. It even infers more plausible trees than obtained via the 100 independent tree inferences conducted with RAxML-NG to identify a best-known ML tree. RAxML-NG is second on all datasets with respect to the number of plausible trees inferred up to a difficulty level of 0.8. On extremely difficult (hopeless) datasets, in analogy to the topological differences, the average number of plausible ML trees inferred with RAxML-NG is roughly on par with the RAxML-NG parsimony starting trees and phylogenies inferred via FastTree2.

The results on simulated data based on RG (using our ad hoc BONK method) data, and the conclusions drawn, are mostly consistent with the results from empirical data in terms of LnL differences and numbers of passed statistical tests. We note, that all tools are able to find trees with higher LnL values than the true tree in some cases. This is not surprising as the ML model is consistent when the number of sites goes to infinity.

For RF- (and Quartet-, see Supplement) distances, however, we observe no substantial differences between RAxML-NG, IQ-TREE2, and FastTree2. This stands in an unexpected contrast to the results on empirical data. We explore this question briefly in the respective Section on differences between simulated and empirical data. Parsimony trees perform consistently worse in terms of RF-distances than all other tools and methods.

Overall, we observe, that for datasets at or above a certain difficulty level (mostly above 0.8), the inference tools become less reliable in reconstructing the true tree and the accuracy differences between various inference tools recede. At these difficulty levels, users should be alarmed and be extremely cautious regarding the subsequent post-analyses and interpretation of the trees. For instance, a small SARS-CoV-2 dataset already exhibits a difficulty of 0.84. Approaches to interpreting and summarizing inference results on such difficult datasets are outlined in Morel *et al*. (42), for instance. We also note, that at least for datasets exhibiting low (easy) and high (hopeless) difficulties, it might be reasonable to reduce the number of independent tree searches conducted with RAxML-NG. This is because the performance differences between the default search on 20 starting trees and the search on 100 distinct starting trees under all metrics deployed here appear to be negligible. In cases, where excessive inference times become an issue, one can also switch to less accurate, yet substantially faster alternatives, such as FastTree2 or even parsimony (e.g., using the fast dedicated parsimony program TNT (43)), without deteriorating the - already low - average inference accuracy.

### Differences between simulated and empirical data

We noticed an unexpected difference between simulated and empirical data in terms of the RF- (and Quartet-) distances between the inferred trees and true trees. There exist several potential explanations for this difference. Apart from the fact that inference and simulation models only represent simplified approximations of real evolutionary processes, we observe, that the value distributions of taxon numbers, number of sites and patterns as well as gap percentages are surprisingly different between TreeBASE and RG (*p* ≪ 0.05 in the two-sample Kolmogorov-Smirnov (KS) test).

Another reason might be our method of gap generation in the simulated MSAs. To assess, whether this constitutes an issue, we conducted an additional set of experiments on simulated data, but this time generated based on TreeBASE meta-data. In contrast to RG, we were able to utilize the original empirical MSA with its corresponding gap patterns. Thus, we first simulated gapless MSAs based on the best-known ML trees (which we also used as reference trees in the empirical tests) and their inferred ML model parameters. Then, we used the gap patterns of the original MSAs and directly superimposed them onto the simulated MSAs. Afterwards, we conducted the same experiments as described above.

This time, both, LnL differences, and topological distances exhibit little variation between the tools, and do not appear to be informative (see Supplement). The observations from the statistical tests are consistent with the previously observed results on simulated RAxMLGrove and empirical TreeBASE data.

Thus, it is less likely that our method for generating realistic gap patterns explains the differences we observe between empirical and simulated datasets. Since we use buckets to group datasets by difficulty, the different difficulty distributions of simulated RG and TreeBASE data (see Figure 6) can also not serve to explain the distinct behavior. Possibly, there is some yet unknown dataset property which we (and the difficulty prediction of Pythia) are not aware of.

**Fig. 5.**
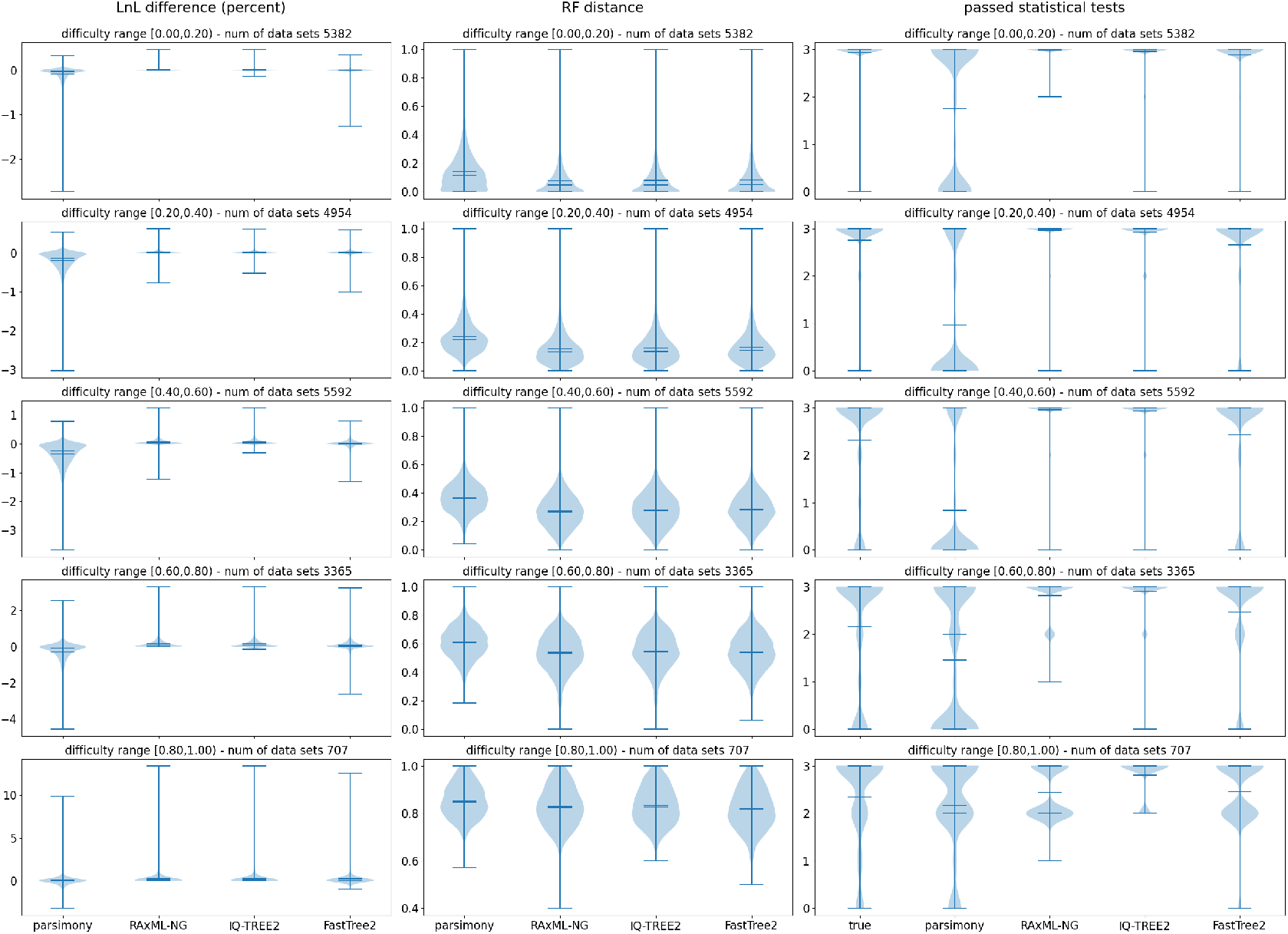
Relative log-likelihood (LnL) score differences, and RF-distances to the true tree, and numbers of passed statistical tests of all inferred trees on simulated MSAs with RAxMLGroveScripts (BONK method).

**Fig. 6.**
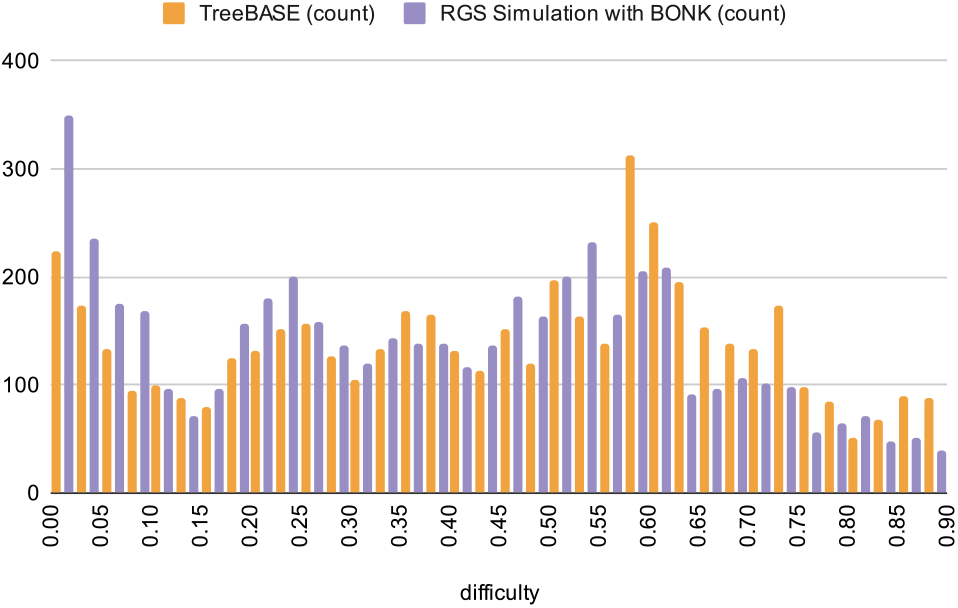
Difficulty distributions of TreeBASE MSAs and simulated RGS MSAs.

Another possible explanation for the discrepancy between results might be a bias towards the more thorough RAxML-NG and IQ-TREE2 tools by defining the “true” tree as the best-known ML tree for the experiments on empirical datasets. We investigated that possibility by conducting an additional experiment on the simulated data. This time, we followed a similar approach as in the empirical data analysis: Out of the set of inferred trees (per dataset), as described before, plus an additional tree obtained via an additional 100 RAxML-NG inferences, we determined the tree with the highest LnL. Then, we computed the RF distances of every inferred tree by the inference tools to the best-known tree based on the LnL score instead of the true tree (see Figure 7). This time, we observe, that the results on simulated data resemble those on empirical data shown in Figure 4. Thus, we confirm that there is a bias introduced by comparing the topology of the inferred trees to the tree with the highest LnL. Note, however, that the RF-distance differences between FastTree2 and RAxML-NG are not as high as in Figure 4. So there are still other not yet understood differences between experiments on empirical and simulated data.

**Fig. 7.**
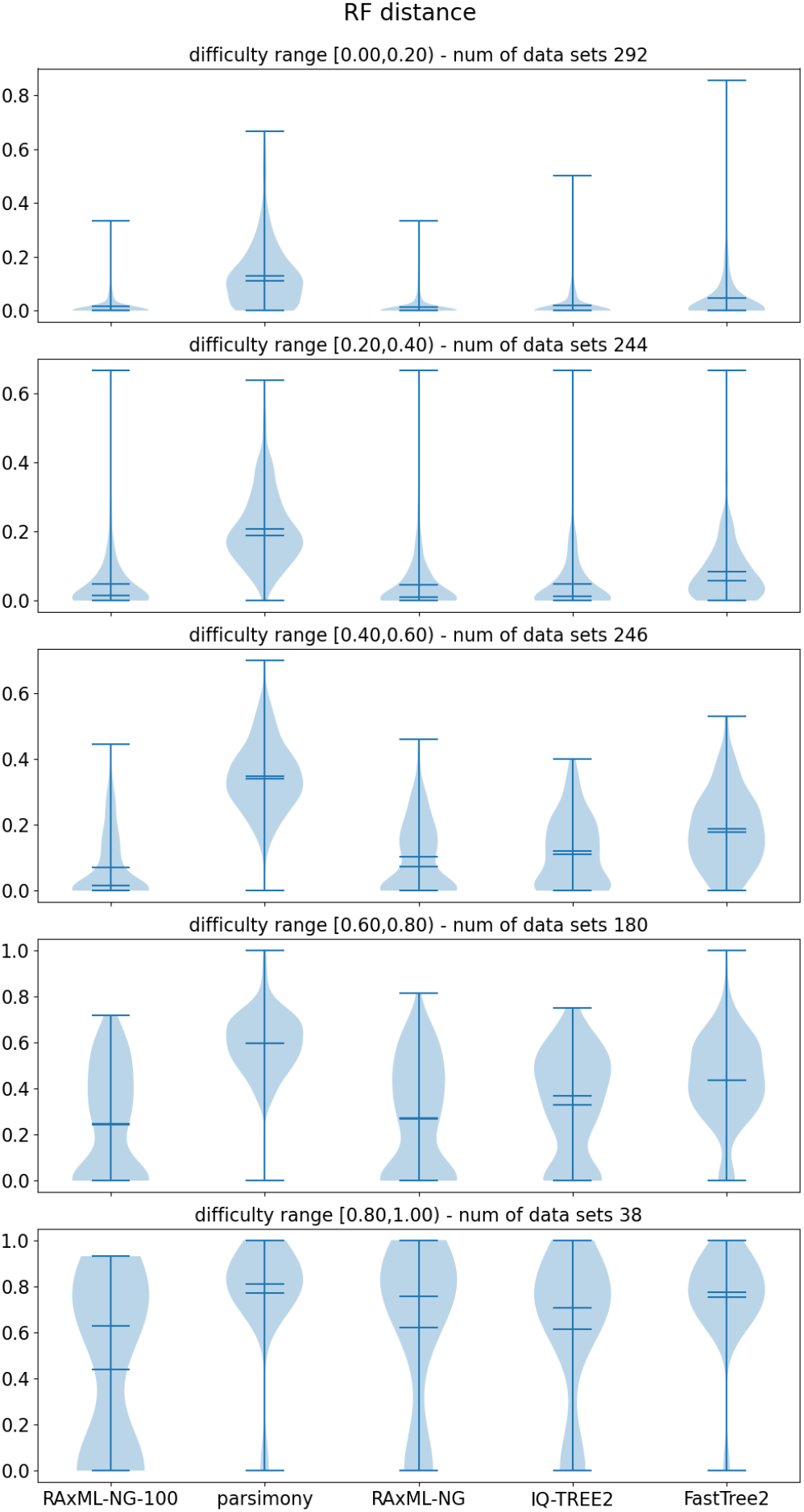
RF differences to best-known ML tree instead of the true tree that we used to simulate the MSAs. “RAxML-NG-100” denotes the tree inferred by conducting 100 tree searches using RAxML-NG.

Overall, we believe, that more research is required regarding the evaluation of tree inference tools. The main focus of ML inference tools is to find trees with the highest LnL score. The main idea of making use of an ML tool, however, is to estimate real evolutionary relationships (the “ground truth”) between species. According to our experiments on simulated data, the ability to find trees with high LnL scores does not necessarily correlate with the ability to infer trees which are topologically close to the ground truth. Therefore, one might question the purpose of running thorough ML optimizations with RAxML-NG or IQ-TREE2 on “common” MSAs instead of inferring a substantially faster (10) evolutionary hypothesis with FastTree2.

## Conclusion

We analyzed the tree inference accuracy of the three widely used phylogenetic inference tools RAxML-NG, IQ-TREE2, and FastTree2 on representative empirical as well as simulated DNA data from TreeBASE and RAxMLGrove respectively. We selected the datasets such that, with respect to their characteristics, they represent the most commonly analyzed datasets by practitioners.

First, we observed a discrepancy between our results on empirical and simulated MSAs, although the simulated MSAs were generated based on empirical parameter distributions. Therefore, more research is required on generating more realistic synthetic datasets that behave like empirical datasets. Second, we observed a bias in the topological tests when inferred trees are compared to the tree with the best LnL score instead of the true tree. When topological reconstruction tests (using the RF distance, for instance) are being conducted on empirical data, one might try to estimate the true tree with the best-known LnL tree, since the true tree (the “ground truth”) is usually unknown. Our experiments on simulated data suggest, that such topological tests can favor the more thorough LnL optimization algorithms. This is an issue to consider when designing and interpreting empirical tests. Since there are no standardized benchmarks, we believe that one should critically assess the evaluation criteria that are typically used for performance assessment. As a first step into this direction, we introduce and make available an easy-to-use Snake-make benchmarking pipeline. This pipeline can help to routinely analyze the accuracy of phylogenetic inference tools in a more standardized manner, under the assumption that Tree-BASE and RAxMLGrove comprise a representative sample of commonly analyzed datasets.

Third, we observed that with increasing difficulty level, as predicted by Pythia, the accuracy of all analyzed tools deteriorates and the differences in accuracy between these tools diminish. This confirms that Pythia implements a meaningful measure for quantifying dataset difficulty in practice. Hence, we recommend applying Pythia before conducting phylogenetic analyses.

Finally, we find that on empirical datasets exhibiting a high difficulty level (difficulty above 0.8), all analyzed tools can essentially be used interchangeably. This means that lengthy computations could – and we would argue *should* – be avoided on such difficult or “hopeless” datasets. More specifically, one should critically assess the necessity of compute- and CO_2_-intensive ML optimization routines, as they are performed by RAxML-NG and IQ-TREE2, especially considering the fact, that – due to possible biases in the evaluation on empirical data – it is not entirely clear, how much more accurate these tools are compared to faster competitors. Therefore, we propose the development of adaptable and flexible heuristic search algorithms that can dynamically take into account the degree of difficulty and the properties of the dataset being analyzed. Hence, the time has come to critically reflect on and rethink the design of heuristic phylogenetic search algorithms, for “What falleth, that shall one also push! […] And him whom ye do not teach to fly, teach I pray you-to fall faster!” (44).

## Acknowledgements

We thank Benoit Morel for helpful discussions, and we thank Antonis Rokas for helpful feedback on this manuscript.

