## Supplement for "A representative Performance Assessment of Maximum Likelihood based Phylogenetic Inference Tools"

### 1 Quartet Distances

Here, we show the quartet distance plots which were omitted from the main manuscript for the sake of readability (see Figures 1, 2, and 3). We also included distances for random trees to present the average ”worst-case” distances.

### 2 AliSim Simulations with Superimposed Gaps

As mentioned in the main manuscript, we conducted an additional experiment on 4994 simulated MSAs using TreeBASE datasets (mentioned in the Section on the differences between simulated and empirical data). We used the AliSim mimick function, which first simulates an MSA, and then superimposes the gap pattern from a reference MSA to the simulated one. The results of these experiments are shown in Figure 4.

### 3 Used Tools

In addition to the tools mentioned in the main manuscript, our implementations of the experimental scripts re-use components from the following libraries and tools: BioPython [1, 6], Google Charts (http://developers.google.com/chart/), ETE 3 [3], Matplotlib [4], Numpy [2], Pandas [5], and SciPy [7].

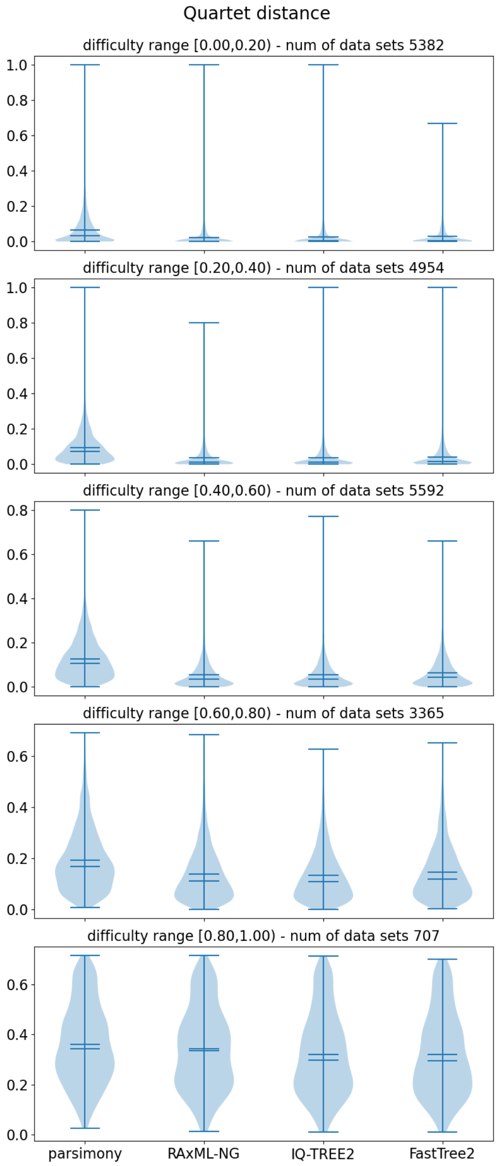


Figure 1: Quartet distances between true trees and inferred trees based on simulated MSAs with RAxMLGroveScripts (BONK method).

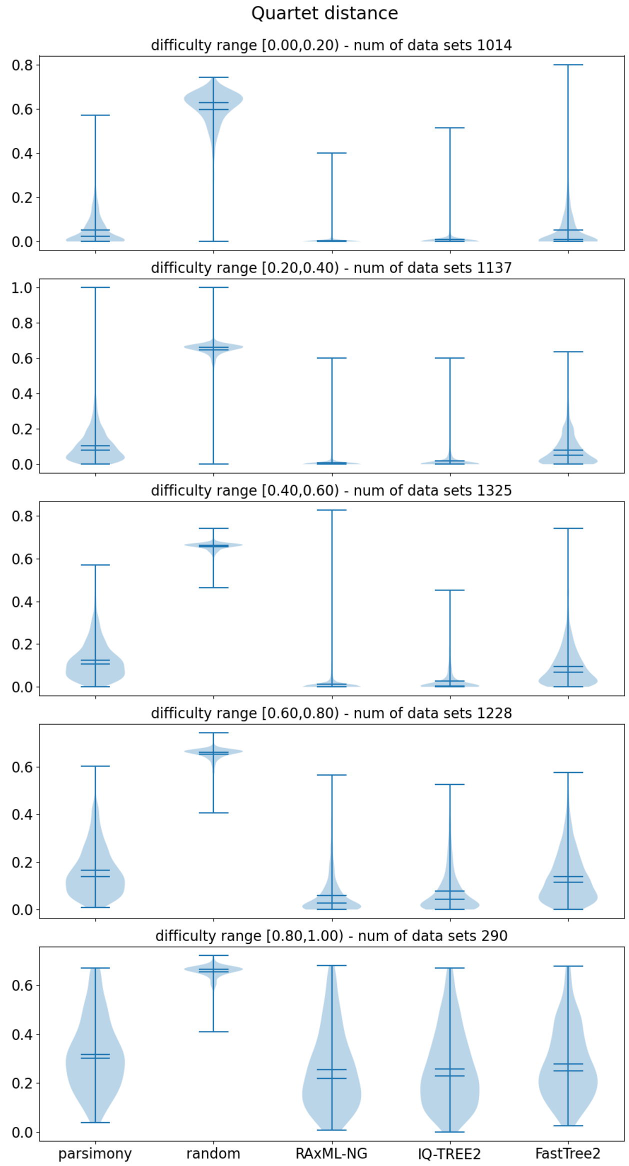


Figure 2: Quartet distances between best known LnL trees and inferred trees based on empirical MSAs from TreeBASE.

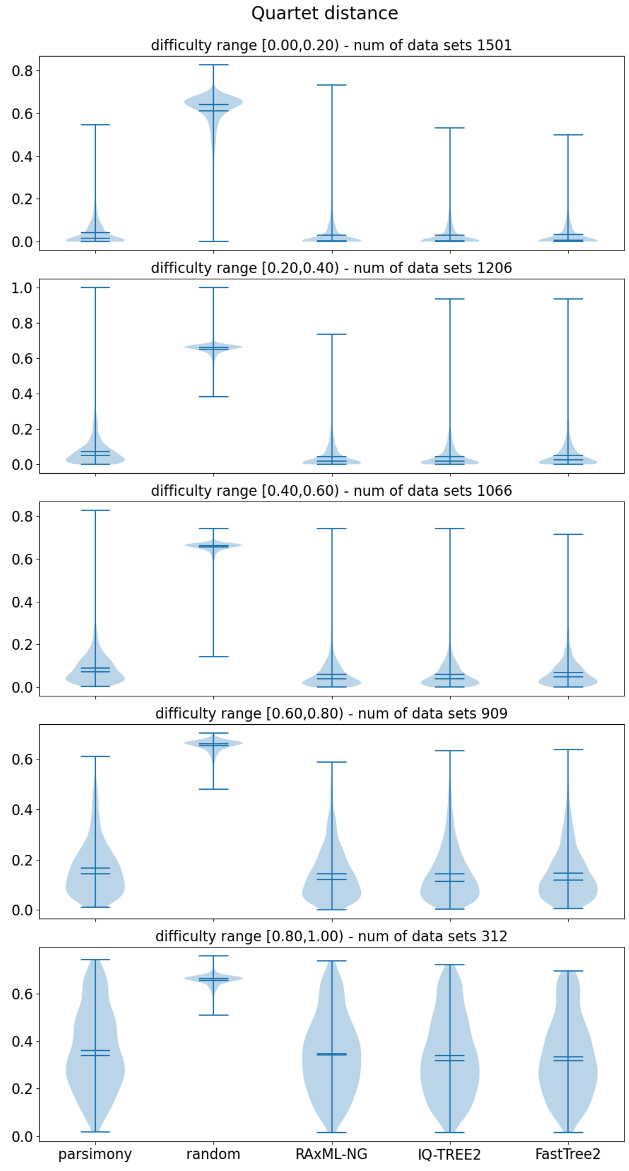


Figure 3: Quartet distances between true trees and inferred trees based on simulated MSAs from TreeBASE data (with superimposed gaps).

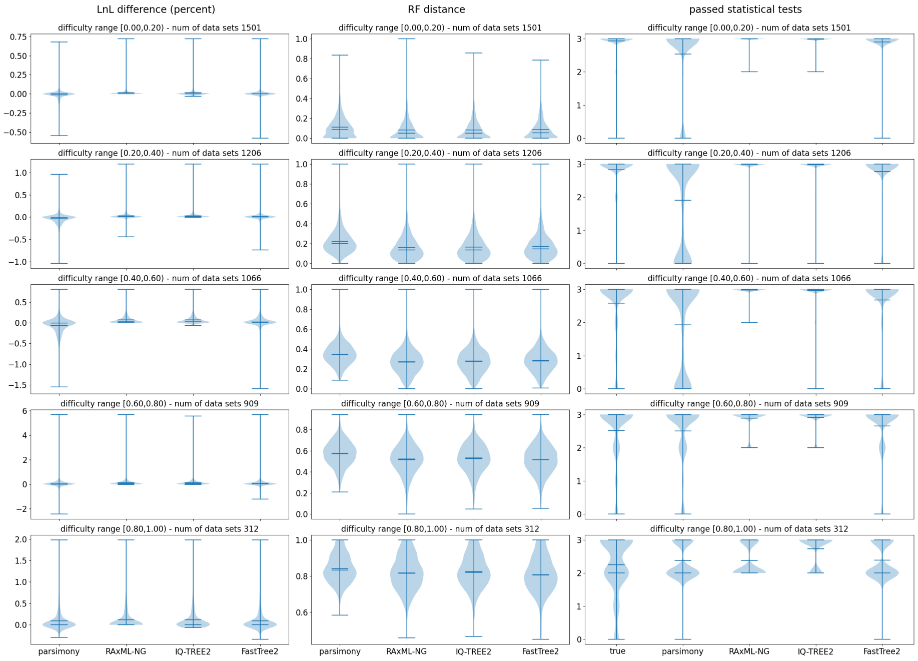


Figure 4: Relative log-likelihood (LnL) score differences, and RF-distances to the true tree, and numbers of passed statistical tests of all inferred trees on simulated MSAs based on TreeBASE data (with superimposed gaps).

[7] Pauli Virtanen, Ralf Gommers, Travis E. Oliphant, Matt Haberland, Tyler Reddy, David Cournapeau, Evgeni Burovski, Pearu Peterson, Warren Weckesser, Jonathan Bright, Stéfan J. van der Walt, Matthew Brett, Joshua Wilson, K. Jarrod Millman, Nikolay Mayorov, Andrew R. J. Nelson, Eric Jones, Robert Kern, Eric Larson, C J Carey, İlhan Polat, Yu Feng, Eric W. Moore, Jake VanderPlas, Denis Laxalde, Josef Perktold, Robert Cimrman, Ian Henriksen, E. A. Quintero, Charles R. Harris, Anne M. Archibald, Antônio H. Ribeiro, Fabian Pedregosa, Paul van Mulbregt, and SciPy 1.0 Contributors. SciPy 1.0: Fundamental Algorithms for Scientific Computing in Python. *Nature Methods*, 17:261–272, 2020. doi: 10.1038/s41592-019-0686-2.
